# Function of *AtPGAP1* in GPI anchor lipid remodeling and transport to the cell surface of GPI-anchored proteins

**DOI:** 10.1101/2021.03.09.434604

**Authors:** César Bernat-Silvestre, Judit Sanchez-Simarro, Yingxuan Ma, Kim Johnson, Fernando Aniento, María Jesús Marcote

**Affiliations:** Departamento de Bioquímica y Biología Molecular, Instituto Universitario de Biotecnología i Biomedicina (BIOTECMED). Universitat de València. Spain; School of BioSciences, University of Melbourne, Parkville, Victoria 3010, Australia; La Trobe Institute for Agriculture & Food, Department of Animal, Plant and Soil Sciences, La Trobe University, Bundoora, Victoria 3086, Australia

**Author notes:** **To whom correspondence should be addressed**, Departamento de Bioquímica y Biología Molecular, Facultad de Farmacia, Universitat de València, Avenida Vicente Andrés Estellés s/n, E-46100 BURJASSOT (VALENCIA). SPAIN. Corresponding author. F.A., K.J. and M.J.M. conceived and designed the experiments. C.B.S., J.S.S and Y.M. performed the experiments. F.A. and M.J.M. wrote the paper, with input from all other authors.

## Abstract

GPI-anchored proteins (GPI-APs) play an important role in a variety of plant biological processes including growth, stress response, morphogenesis, signalling and cell wall biosynthesis. The GPI-anchor contains a lipid-linked glycan backbone that is synthesized in the endoplasmic reticulum (ER) where it is subsequently transferred to the C-terminus of proteins containing a GPI signal peptide by a GPI transamidase. Once the GPI anchor is attached to the protein, the glycan and lipid moieties are remodelled. In mammals and yeast, this remodelling is required for GPI-APs to be included in Coat Protein II (COPII) coated vesicles for their ER export and subsequent transport to the cell surface. The first reaction of lipid remodelling is the removal of the acyl chain from the inositol group by Bst1p (yeast) and PGAP1 (mammals). In this work, we have used a loss-of-function approach to study the role of *PGAP1/Bst1* like genes in plants. We have found that *Arabidopsis* PGAP1 localizes to the ER and probably functions as the GPI inositol-deacylase which cleaves the acyl chain from the inositol ring of the GPI anchor. In addition, we show that PGAP1 function is required for efficient ER export and transport to the cell surface of GPI-APs.

**One sentence summary:** GPI anchor lipid remodeling in GPI-anchored proteins is required for their transport to the cell surface in *Arabidopsis*.

## INTRODUCTION

Proteins associated with the plasma membrane are involved in a variety of essential functions in eukaryotes, including signalling, transport and cell surface metabolism (Yeats et al., 2018). Proteins can be attached to the plasma membrane several ways. Transmembrane proteins contain domains with hydrophobic amino acids which are embedded within the plasma membrane lipid bilayer, while other proteins use a post-translational attachment to lipids. For instance, if a protein has to be on the intracellular face of the plasma membrane, it can be post-translationally modified by *S*-acylation, *N*-myristoylation, prenylation or palmitoylation (Luschnig and Seifert, 2011; Hemsley, 2015). A protein can also be attached to a GPI anchor during secretion, which targets it to the outer surface of the plasma membrane.

GPI-anchored proteins (GPI-APs) have been studied from yeast and trypanosomes to mammals and plants and shown to be involved in crucial biological processes, including growth, morphogenesis, reproduction and pathogenesis (Cheung et al., 2014). The GPI anchor is newly synthesized in the ER and is then attached to the C-terminus of proteins containing a GPI signal peptide by a GPI transamidase (Desnoyer et al., 2020; Kinoshita, 2020). The GPI anchor precursor consists of a glycan core (with glycan side chains) and phosphatidylinositol (PI) with an acyl chain at the 2 position of the inositol ring. The glycan core is characterized by having a N-acetyl glucosamine and three α-linked mannoses attached to ethanolamine phosphate (EtNP). The GPI anchor is attached to the polypeptide by an amide bond between EtNP and the C-terminal of the polypeptide (Kinoshita and Fujita, 2016). The GPI-AP stays anchored to the luminal leaflet of the ER lipid bilayer by insertion of hydrocarbon chains of PI, diacyl-PI in yeast and a mixture of 1-alkyl-2-acyl PI (major form) and diacyl-PI (minor form) in mammalian cells. Once the GPI anchor is attached to the protein, the glycan and lipid moieties are remodelled and this process has been shown to be very important for ER export and transport to the cell surface of GPI-APs (Muñiz and Riezman, 2016; Kinoshita, 2020).

This remodelling has been extensively studied in yeast and mammals. In yeast, the GPI-lipid remodelling occurs entirely in the ER (Pittet and Conzelmann, 2007) and it is initiated with the inositol deacylation at the 2 position of the inositol ring by the remodelling enzyme Bst1p (PGAP1 in mammals) (Figure 1) (Tanaka et al., 2004; Fujita et al., 2006b). This makes GPI-APs sensitive to the bacterial phosphatidylinositol-specific phospholipase C (PI-PLC) (Low, 1989). Next, the short and unsaturated fatty acid (C18:1) at the sn2 position of PI is removed by the remodelling enzyme Per1p (Figure 1) (Fujita et al., 2006a) and it is replaced with a very long-chain saturated fatty acid (C26:0) by Gup1p (Bosson et al., 2006). The C26:0 DAG generated seems to be present only in those GPI-APs destined to be transferred to the cell wall (Pittet and Conzelmann, 2007). GPI-APs that remain in the plasma membrane also contain anchors with a very long-chain saturated fatty acid (C26:0) in the form of ceramide. Cwh43p is the enzyme which carries out the addition of ceramide as the lipid moiety to the GPI anchor (Umemura et al., 2007; Yoko-o et al., 2018). The substrate for the ceramide remodelling is still not clear, but it has been described that most lipid moieties of GPI anchors are exchanged from diacylglycerol to ceramide types (Ghugtyal et al., 2007). These long-chain saturated fatty acids change the physical properties of the GPI-APs and the association with the membrane forming ordered domains at the ER lipid membrane (Silva et al., 2006) allowing these domains to be selectively concentrated at specific ER exit sites (ERES) (Muñiz and Riezman, 2016; Rodriguez-Gallardo et al., 2020). The GPI-APs preassembled at ERESs are transported from the ER to the Golgi by Coat Protein II (COPII) coated vesicles. Due to the luminal localization of GPI-APs, a cargo receptor (p24 complex) is required for incorporation of GPI-APs into nascent COPII vesicles (Castillon et al., 2011; Manzano-Lopez et al., 2015). Then, GPI-APs are transported along the secretory pathway, through the Golgi complex, to their final destination, the plasma membrane or the cell wall. Proper fatty acid remodelling of the GPI anchor allows GPI-APs to associate to membrane microdomains enriched in sphingolipids and cholesterol (lipid rafts) (Fujita et al., 2006a; Maeda et al., 2007; Castillon et al., 2011; Muniz and Zurzolo, 2014).

**Figure 1.**
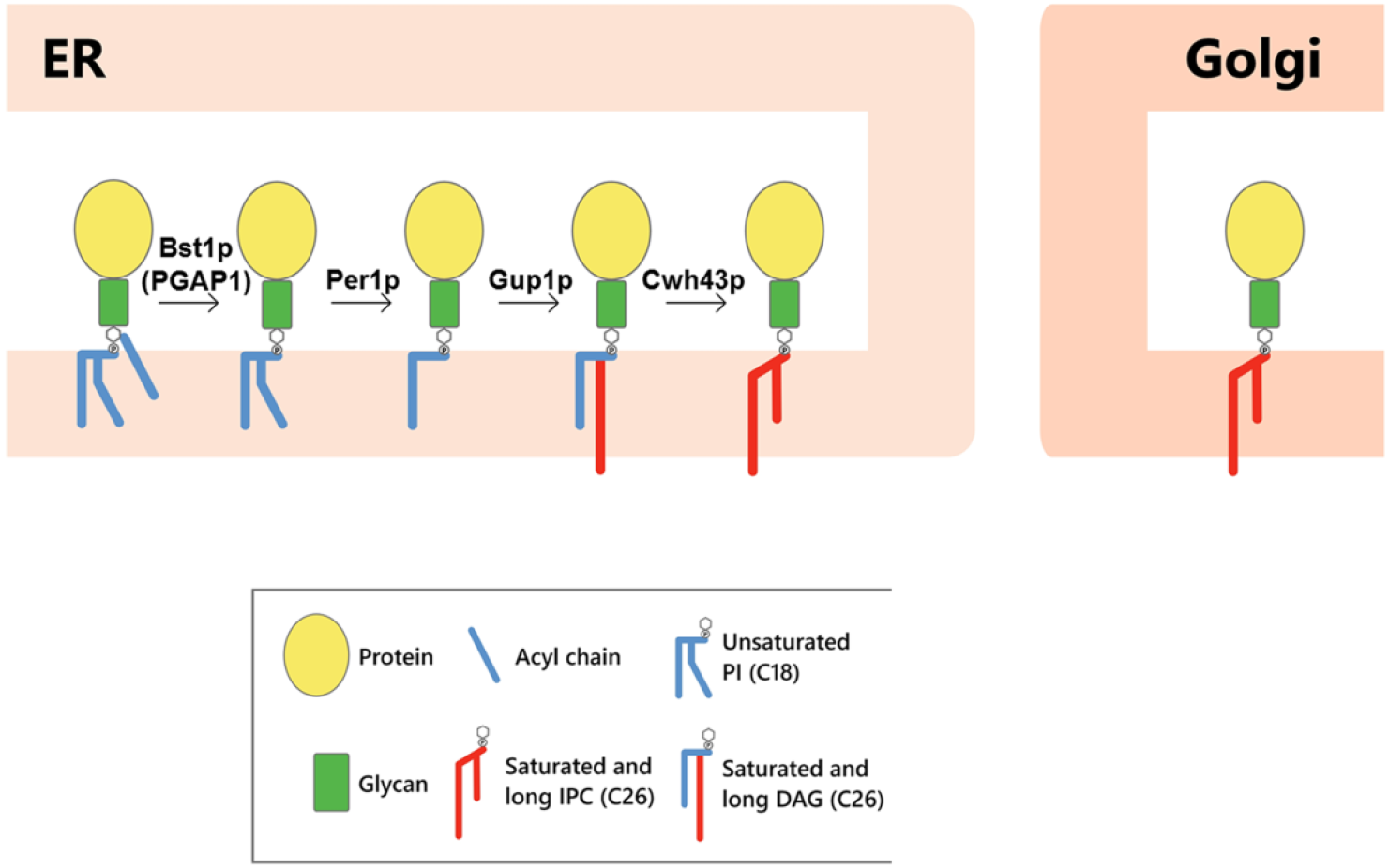
GPI-lipid remodeling in yeast. The GPI anchor is synthesized in the ER and consists of a glycan core and diacyl-phosphatidylinositol (PI) with an acyl chain at the 2 position of the inositol ring (Kinoshita and Fujita, 2016). After protein attachment, the glycan and lipid moieties are remodelled and this process has been shown to be very important for the transport and final localization of the GPI-APs. The GPI-lipid undergoes a structural remodeling that has the purpose of providing saturated lipids and in yeast it occurs almost entirely at the ER where it is initiated with the inositol deacylation at the 2 position of the inositol ring by the remodelling enzyme Bst1p (PGAP1 in mammals). The rest of lipid remodeling enzymes are indicated, see text for details (modified from Muniz and Zurzolo, (2014)).

In *Arabidopsis thaliana* it has been predicted around 300 GPI-APs (10 % of secreted proteins) and they play an important role in a variety of plant biological processes occurring at the interface of the plasma membrane and the cell wall, including growth, stress response, morphogenesis, signalling, cell wall biosynthesis and maintenance or plasmodesmatal transport (Borner et al., 2003; Yeats et al., 2018). Up to now, only one plant GPI anchor structure has been resolved, the structure of *Pc*AGP1, isolated from *Pyrus communis* (pear) cell suspension culture (Oxley and Bacic, 1999). From this structure, it can be said that the core structure of GPI anchors seems to be conserved in plant and non-plant eukaryotes. In addition, a survey of the *Arabidopsis* genome indicates that most of the genes involved in particular steps of GPI anchor assembly and their remodeling have orthologs in *Arabidopsis* (Luschnig and Seifert, 2011). Disrupting GPI-anchor synthesis in *Arabidopsis* is lethal, as is the case in yeast and mammals (Lalanne et al., 2004; Gillmor et al., 2005; Dai et al., 2014; Bundy et al., 2016). However, no studies have been reported of lipid remodelling enzymes of GPI-APs in plants. Such studies would provide important insights to understand plant GPI-anchor remodelling and its role in trafficking and function of GPI-APs, as it was the case with studies in mammals and yeast (Muñiz and Riezman, 2016; Kinoshita, 2020), and may reveal plant distinct and unique characteristics. In this work, we have used a loss-of-function approach to initiate the study of the role of an *Arabidopsis* ortholog of mammalian PGAP1 and yeast Bst1p, the enzymes involved in inositol deacylation, the first step of GPI anchor remodelling.

## RESULTS

### 1. Identification of PGAP1 in *Arabidopsis*

Inositol deacylation of GPI-APs is mediated by mammalian PGAP1 and yeast Bst1p (Figure 1). Yeast Bst1p (1029 amino acids) and mammalian PGAP1 (922 amino acids) are multi transmembrane ER proteins with a catalytic serine containing motif that is conserved in a number of lipases (Tanaka et al., 2004). By searching for *Arabidopsis* putative GPI inositol-deacylase PGAP1-like (IPR012908, pfam07819) genes using Pfam and InterPro databases (Hunter et al., 2009; Finn et al., 2010), 6 *Arabidopsis* genes have been found (Supplemental Table 1). In this study, AT3G27325 was chosen for further investigation, as it is the only *Arabidopsis PGAP1-like* gene that encodes a putative ER transmembrane protein (Supplemental Table 1) (Luschnig and Seifert, 2011). As shown in Supplemental Figure 1, the lipase consensus sequence is conserved between AT3G27325, hsPGAP1 and Bst1p. In this study, we chose *PGAP1* (the mammalian name of the gene) to name AT3G27325 because it is the name that appears in the *Arabidopsis* Information Resource (TAIR) and because the yeast name (Bst1) had already been assigned to the *Arabidopsis* gene AT5G65090 (*BST1*, Bristled). To investigate the expression of *PGAP1* in different developmental stages, we used the publicly available RNAseq expression database GENEVESTIGATOR (www.genevestigator.com) (Zimmermann et al., 2004; Hruz et al., 2008). PGAP1 shows medium levels of expression in most tissues throughout plant development (Supplemental Figure 2).

In order to determine the subcellular localization of PGAP1, a construct of PGAP1 with C-terminal RFP was used for transient expression in *N. benthamiana* leaves. PGAP1-RFP showed an ER-like localization pattern and extensively colocalized with the ER marker GFP-HDEL (Figure 2). No colocalization was found with markers of the Golgi apparatus (YFP-ManI) or ER export sites (ERES)/cytosol (GFP-SEC24C). These results clearly showed that Arabidopsis PGAP1 protein localizes to the ER, consistent with the localization of mammalian PGAP1 (Tanaka et al., 2004) and yeast Bst1p (Elrod-Erickson and Kaiser, 1996).

**Figure 2.**
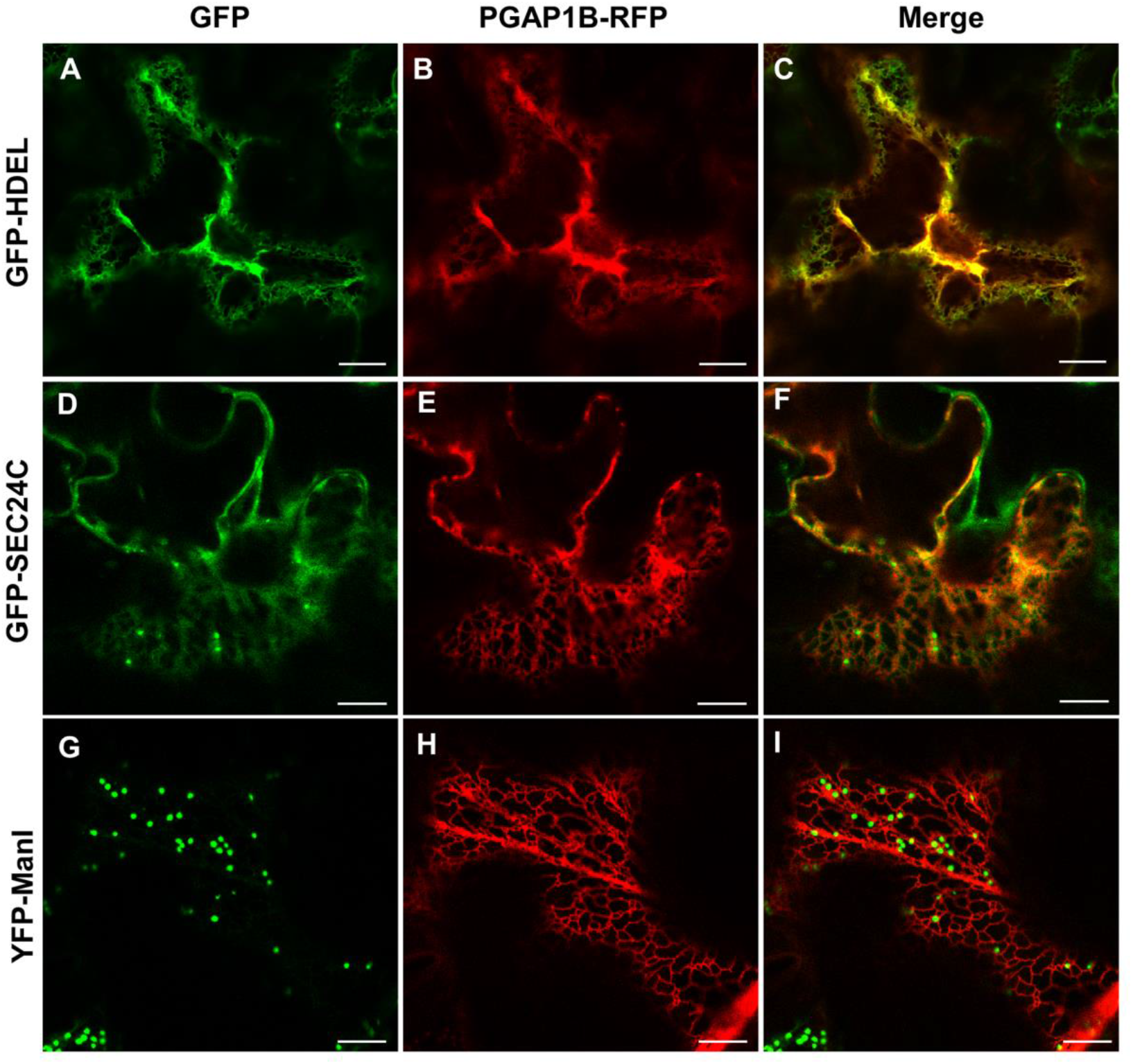
Localization of PGAP1-RFP. Transient gene expression in *N. benthamiana* leaves of PGAP1-RFP (B, E, H) together with GFP-HDEL (ER marker) (A), GFP-SEC24C (ERES and cytosol marker) (D) and YFP-ManI (Golgi marker) (G) (see merged images in C, F and I). Scale bars = 10 μm.

To functionally characterize *Arabidopsis PGAP1*, a reverse genetics approach was chosen. Several T-DNA insertion mutants of *PGAP1* were found in the *Arabidopsis* SALK collection (http://signal.salk.edu/cgi-bin/tdnaexpress). Four mutants of *PGAP1, pgap1-1* (SALK_078662), *pgap1-2* (SAIL_1212_H07), *pgap1-3* (Salk_027086) and *pgap1-4* (Salk_004218), with T-DNA insertions in different positions within the gene, were characterized (Figure 3). Homozygous plants were selected by PCR and RT-PCR analysis showed that all lines were knock-down mutants (Figure 3). All *pgap* mutants showed less than 10% of wild-type *PGAP1* expression levels. We selected *pgap1-1* and *pgap1-3* mutants for further analysis. *pgap1* mutants showed slightly reduced plant height compared to wild-type plants with no other obvious phenotypic differences under standard growth conditions (Figure 3). GPI-APs in plants have been shown to influence cell wall metabolism, cross-linking and signaling (reviewed in (Yeats et al., 2018)). To investigate possible changes in cell walls in *pgap1* mutants, cell wall compositional analysis was undertaken. Analysis of cell wall polysaccharides showed increased amounts of arabinan and decreased content of type II AG and Xyloglucan in the *pgap1-3* mutant compared to wild-type (Figure 4 and Supplemental Table 2).

**Figure 3.**
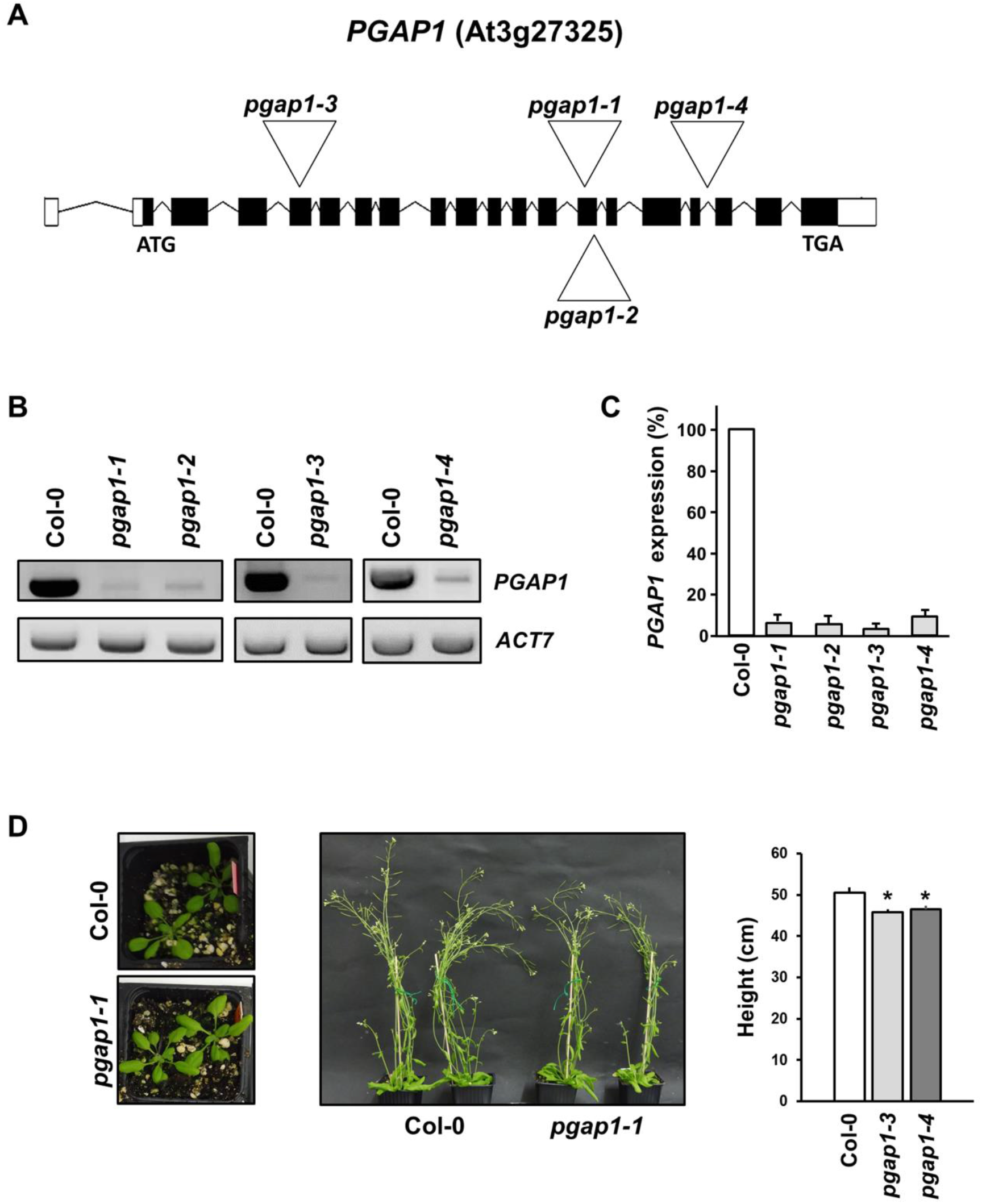
Characterization of *pgap1* mutants. **A.** Diagram of the *PGAP1* gene and localization of the T-DNA insertion (triangle) in the *pgap1-1* and *pgap1-2, pgap1-3* and *pgap1-4* mutants. Black boxes represent coding regions and white boxes represent 5’ UTR and 3’ UTR regions. **B.** sqPCR analysis of *PGAP1* expression in *pgap1* mutants. Total RNA from *pgap1-1, pgap1-2, pgap1-3*, *pgap1-4* and wild-type (Col-0) 4 day-old seedlings were used for the PCR. In the PCRs, *PGAP1* specific primers were used (Supplemental Table 4). *Actin-7 (ACT7)* was used as a control. PCR samples were collected at cycle 22 for *ACT7* and at cycle 36 for *PGAP1*.**C** Quantification of the experiments shown in B from three biological samples. Values were normalized against the *PGAP1* fragment band intensity in wild type that was considered to be 100%. Error bars represent SEM. All the four mutants were knock-down mutants as a fragment of the cDNA of *PGAP1* was detected in all of them. **D** Left panel, 20 day -old plants and middle panel, 42 day-old plants of wild-type and the *pgap1* mutants, respectively. In the right panel, the height of 42 day-old wild-type and *pgap1* mutant plants expressed as mean ± s.e.m (n=4). Data Asterisk represents statistic differences based on Student’s t test with P<0.05.

**Figure 4.**
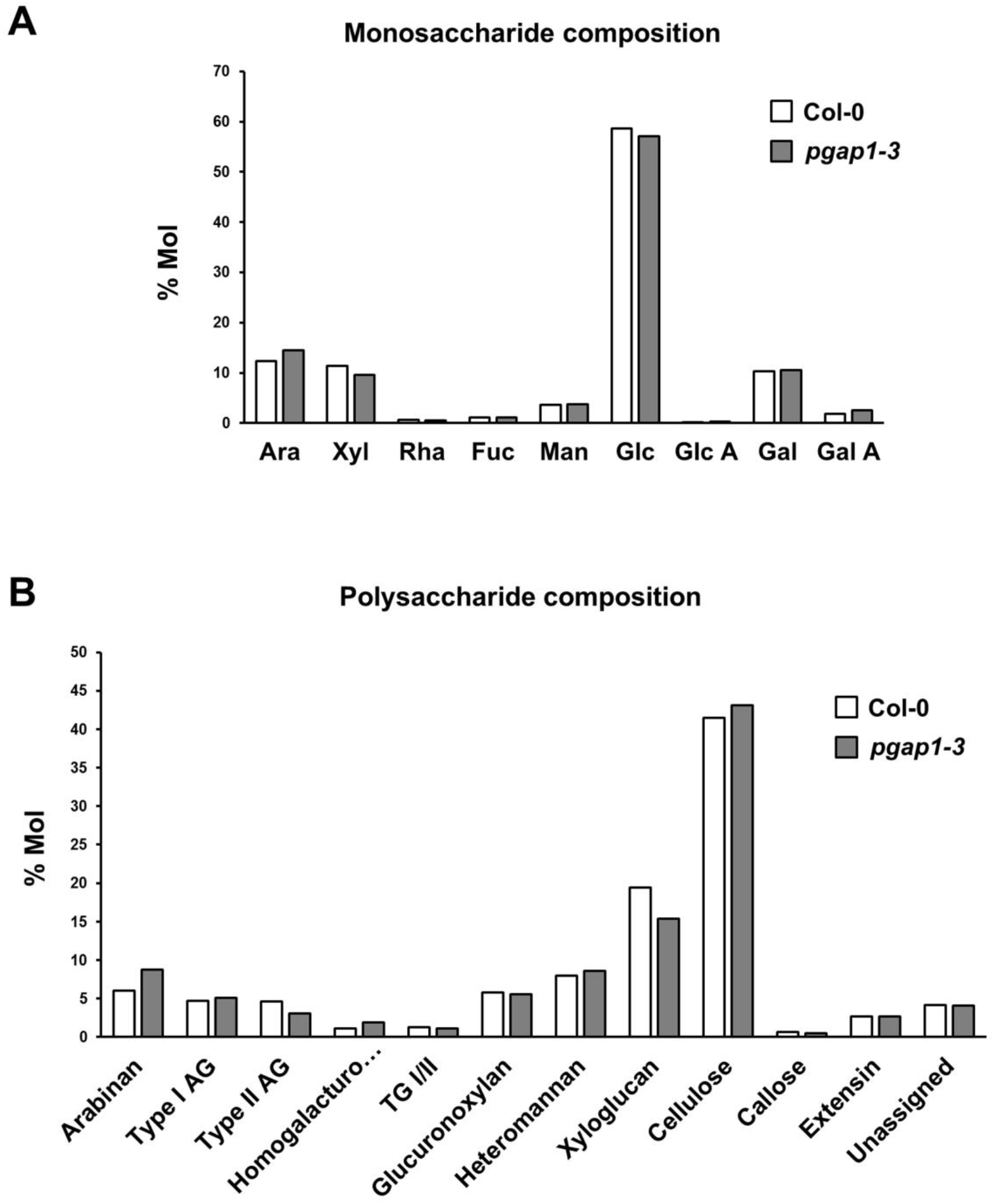
Monosaccharide and polysaccharide compositions of wild-type (Col-0) and *pgap1-3* seedlings. Methylation analysis showed no obvious differences in deduced monosaccharide (A) composition between Col-0 and *pgap1-3* mutants. Polysaccharide composition (B) showed a trend of reduced Type II AG and Xyloglucan and increased Arabinan content in *pgap1-3* compared to Col-0. Ara: arabinose. Xyl: xylose. Rha: rhamnose. Fuc: fucose. Man: mannose. Glc: glucose. GlcA: glucuronic acid. Gal: galactose. GalA: galacturonic acid. AG: arabinogalactan. RG: rhamnogalacturonan. Data shown are the average of two biological replicates with two technical replicates each.

### 2. Localization of GPI-anchored proteins in *pgap1* mutants

GPI anchor remodeling has been shown to be important for efficient trafficking of yeast and mammalian GPI-anchored proteins from the ER to the plasma membrane (Tanaka et al., 2004; Fujita et al., 2006b; Castillon et al., 2009; Castillon et al., 2011). Therefore, we analyzed the localization of GPI-anchored proteins in *pgap1* mutants. To this end, we used two different GPI-APs. One of them was GFP fused to arabinogalactan protein 4 (AGP4). Arabinogalactan proteins (AGPs) are cell surface proteoglycans that seem to be involved in diverse developmental processes such as differentiation, cell-cell recognition, embryogenesis and programmed cell death (Ellis et al., 2010; Pereira et al., 2016; Strasser et al., 2021). GFP-AGP4 has been shown previously to localize to the plasma membrane (Martinière et al., 2012; Bernat-Silvestre et al., 2020). The second one was Venus fused to FLA11 (V-FLA11), a member of Fasciclin-Like Arabinogalactan proteins (FLAs) that play important biological roles related to cell adhesion (Johnson, 2003; MacMillan et al., 2010). In addition, we also used a glycosylphosphatidylinositol-anchored GFP (GFP-GPI) (Martinière et al., 2012; Bernat-Silvestre et al., 2020). As a control, we used a transmembrane plasma membrane protein, the aquaporin PIP2A-RFP (Nelson et al., 2007).

We first analyzed the localization of these proteins by transient expression in *Arabidopsis* seedlings. As shown in Figure 5, GFP-AGP4, V-FLA11 and GFP-GPI were exclusively localized to the plasma membrane of cotyledon cells of wild-type *Arabidopsis* seedlings, as was the case for the transmembrane plasma membrane protein PIP2A-RFP. This was shown previously for GFP-AGP4 and GFP-GPI (Bernat-Silvestre et al., 2020). V-FLA11 was also found to colocalize with PIP2A-RFP in wild-type seedlings (Supplemental Figure 3). In clear contrast, GFP-AGP4, V-FLA11 and GFP-GPI showed an ER-like localization pattern in *pgap1* mutants, which was not the case of PIP2A-RFP, which localized to the plasma membrane in these mutants. This suggests that PGAP1 enzyme is specifically required for transport to the plasma membrane of GPI-anchored proteins, and that loss of *PGAP1* function does not affect transport from the ER to the plasma membrane of transmembrane proteins. The defect in transport of GPI-APs in *pgap1* mutants was not a consequence of an alteration in the compartments of the secretory pathway, since no obvious defects were observed in the localization pattern of several organelle marker proteins, including GFP-HDEL (ER), GFP-EMP12 (Golgi apparatus), TIP1-GFP (tonoplast), SPdCt-mCherry (vacuole lumen), SCAMP1-YFP (plasma membrane) and GFP-CESA3 (TGN/plasma membrane) (Supplemental Figure 4). The ER localization of GFP-AGP4 and V-FLA11 in *pgap1* mutants was then confirmed by colocalization experiments. As shown in Figure 6, both GFP-AGP4 and V-FLA11 strongly colocalized with two different ER marker proteins, an ER luminal protein (mCherry-HDEL) and an ER membrane protein (RFP-p24δ5). Occasionally, GFP-AGP4 and V-FLA11 were also found in punctate structures, which colocalized with the Golgi marker RFP-ManI, suggesting that a small fraction of these GPI-anchored proteins also localize to the Golgi apparatus in *pgap1* mutants. The main ER localization of GFP-AGP4 and V-FLA11 in *pgap1* mutant *Arabidopsis* seedlings was also confirmed biochemically (see below, Figure 8A).

**Figure 5.**
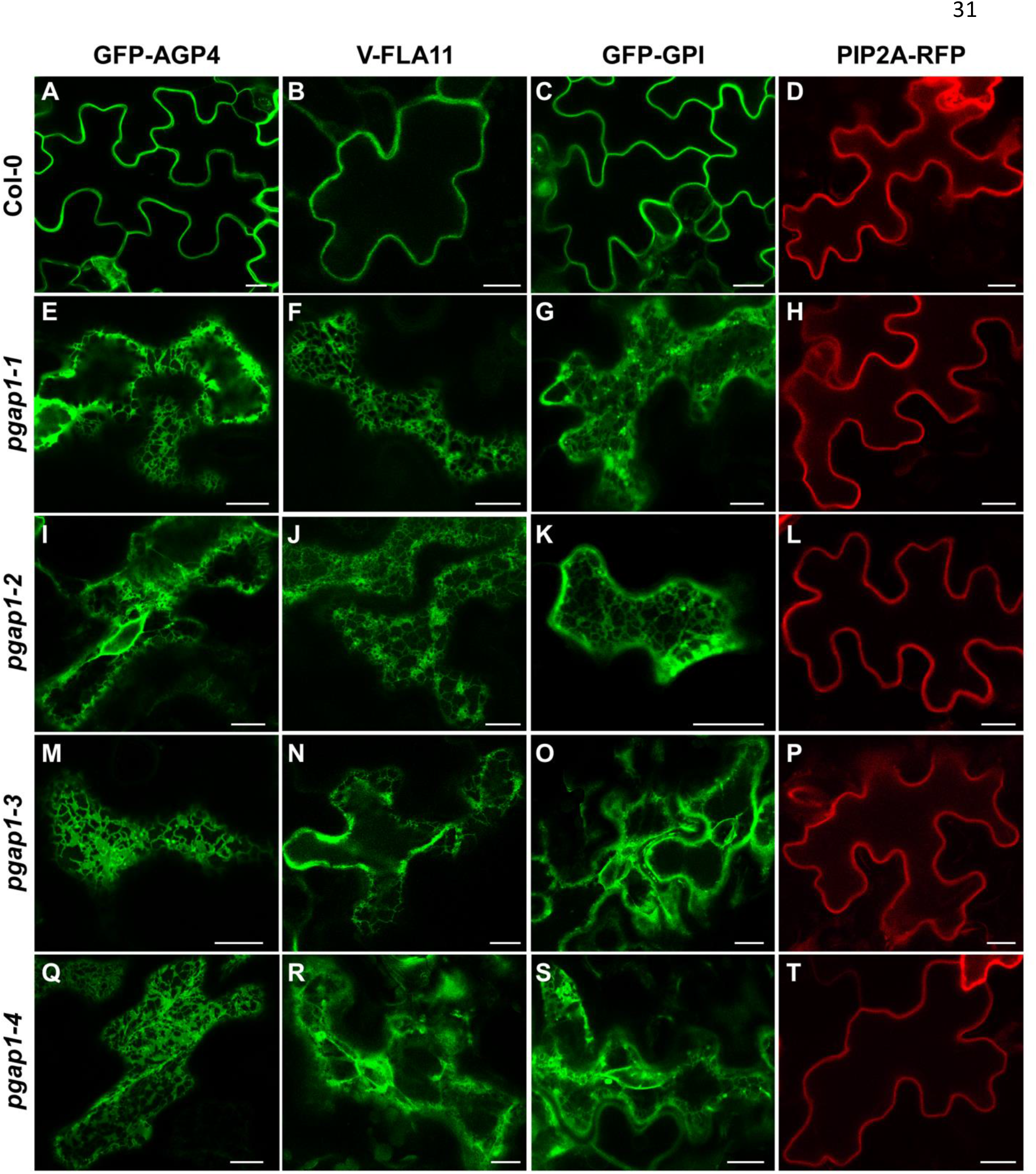
Localization of GFP-AGP4, V-FLA11 and GFP-GPI in wild-type and *pgap1* mutant *Arabidopsis* seedlings. Transient gene expression in wild-type (Col-0) (A-D) or *pgap1-1* (E-H), *pgap1-2* (I-L), *pgap1-3* (M-P) and *pgap1-4* (Q-T) *Arabidopsis* seedlings. The three GPI-anchored proteins, GFP-AGP4, V-FLA11 and GFP-GPI mainly localized to the plasma membrane in cotyledon cells from wild-type (Col-0) seedlings, as the transmembrane plasma membrane protein PIP2A-RFP. In the 4 *pgap1* mutants, GFP-AGP4, V-FLA11 and GFP-GPI showed a predominant ER localization pattern, in contrast to PIP2A-RFP, which mainly localized to the plasma membrane. Scale bars, 10 μm.

**Figure 6.**
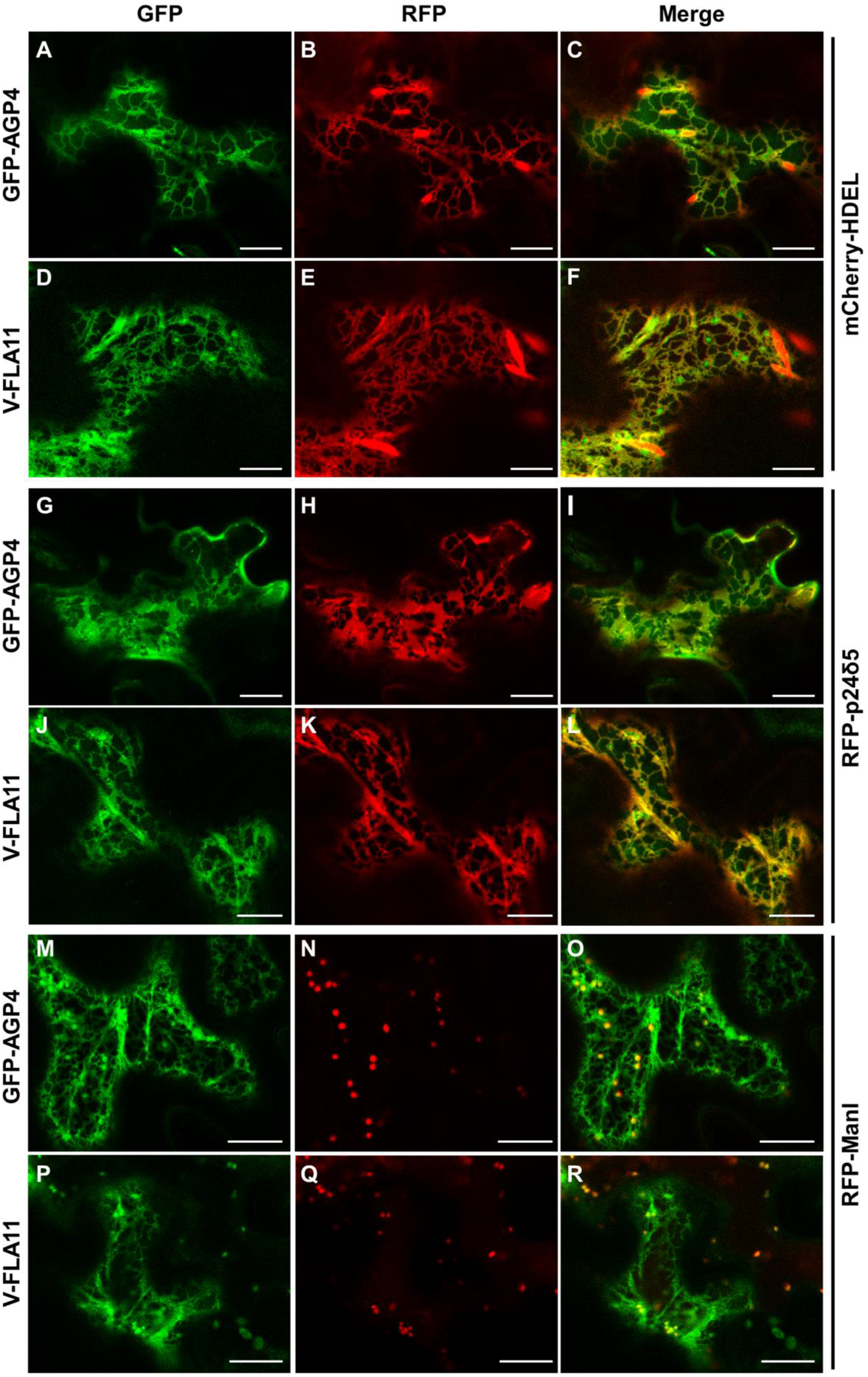
Colocalization of GPI-APs with ER markers in *pgap1-3* mutant seedlings. Transient gene expression in *Arabidopsis* seedlings. **(A-F)**. Coexpression of GFP-AGP4 (A) and V-FLA11 (D) with the ER marker mCherry-HDEL (B, E) (see merged images in C and F). **(G-L)**. Coexpression of GFP-AGP4 (G) and V-FLA11 (J) with the ER marker RFP-p24δ5 (H, K) (see merged images in I and L). **(M-R)**. Coexpression of GFP-AGP4 (M) and V-FLA11 (P) with the Golgi marker RFP-ManI (N, Q) (see merged images in O and R). Scale bars = 10 μm.

**Figure 7.**
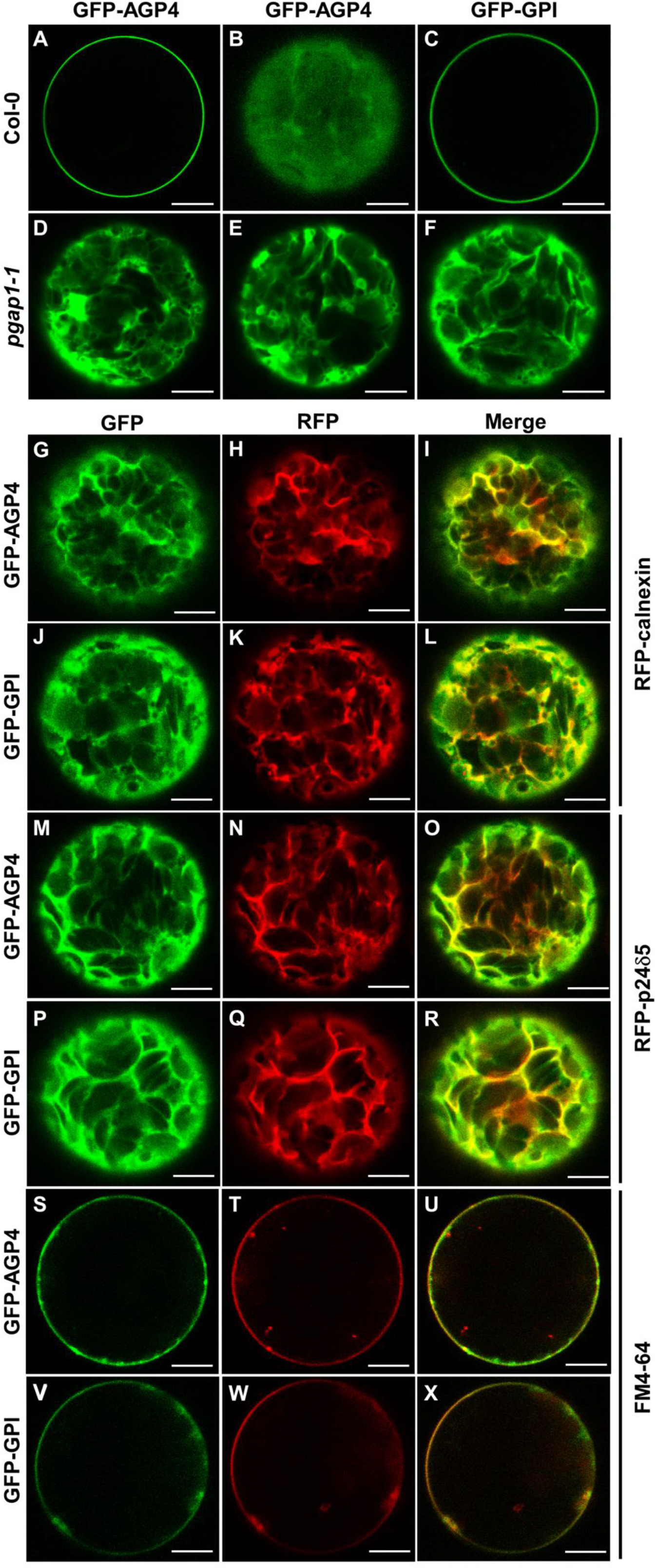
Localization of GFP-AGP4 and GFP-GPI in wild-type and *pgap1-1 Arabidopsis* protoplasts. Transient expression in wild-type (A-C) and *pgap1-1 Arabidopsis* protoplasts (D-X). **(A-F)** GFP-AGP4 (A, B) and GFP-GPI (C) localized in the plasma membrane in wild-type protoplasts. In contrast, GFP-AGP4 (D, E) and GFP-GPI (F) showed an ER-like localization pattern in *pgap1-1* protoplasts (D-F). **(G-L)**. Coexpression of GFP-AGP4 (G) or GFP-GPI (J) with the ER marker RFP-calnexin (H, K) (see merged images in I and L). **(M-R)**. Coexpression of GFP-AGP4 (M) or GFP-GPI (P) with the ER marker RFP-p24δ5 (N, Q) (see merged images in O and R). **(S-X)**. Colocalization of GFP-AGP4 (S) or GFP-GPI (V) with the FM dye FM4-64 (T, W) (see merged images in U and X).

**Figure 8.**
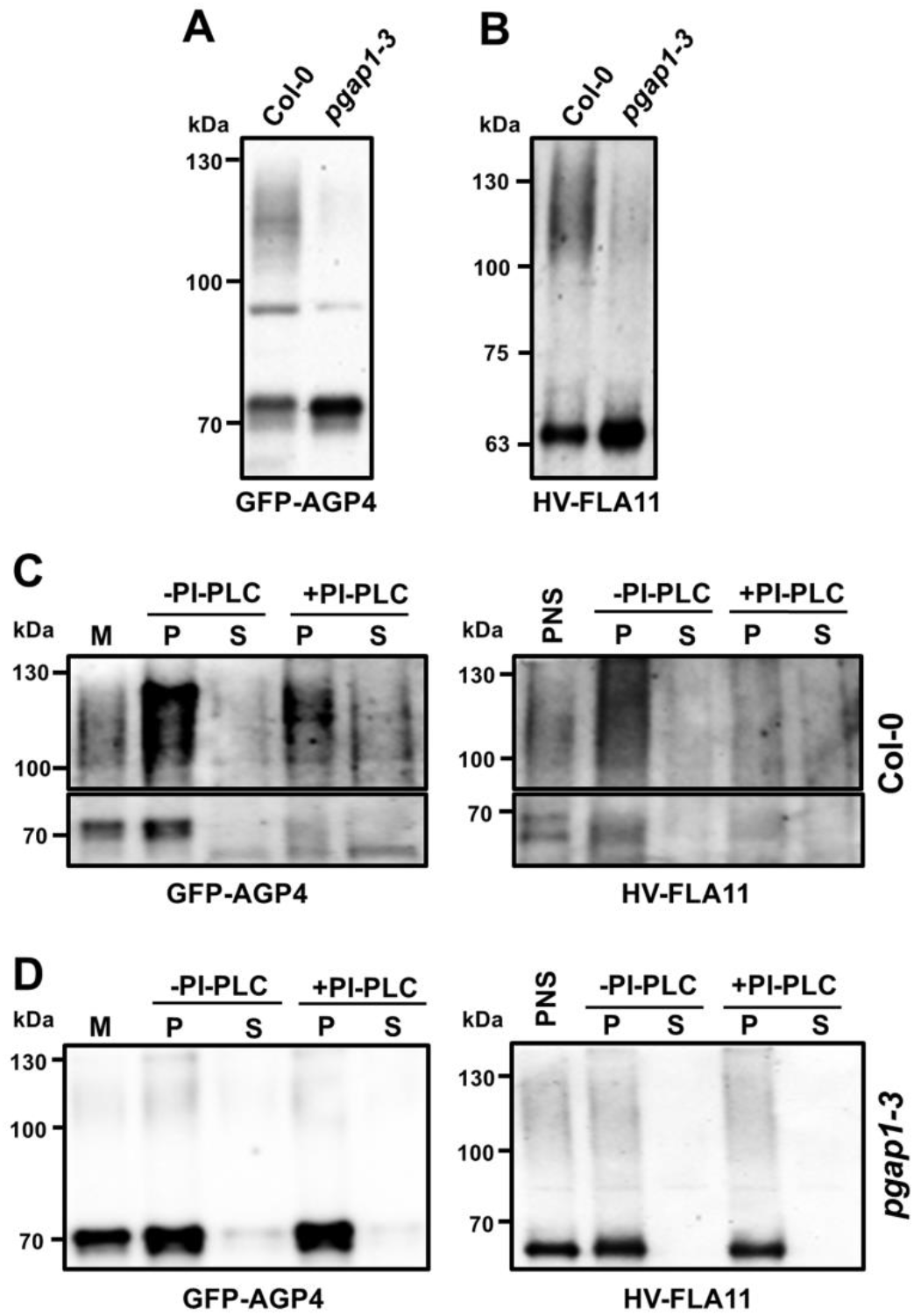
Biochemical characterization of GFP-AGP4 and V-FLA11 in wild-type and *pgap1-3* mutant seedlings. **A-B.** Postnuclear supernatants (PNS) were obtained from cotyledons of wild-type (Col) and *pgap1-3* mutant seedlings transiently expressing GFP-AGP4 (A) and V-FLA11 (B) and analyzed by SDS-PAGE and immunoblotting with antibodies against GFP (to detect GFP-AGP4 and V-FLA11). In wild-type seedlings, both GFP-AGP4 and V-FLA11 showed a smear with a molecular mass around 100-130 kDa, which correspond to the plasma membrane form of GFP-AGP4 (Bernat-Silvestre, 2021) and V-FLA11 (Supplemental Figure 8), and additional bands (around 70 kD and 63 kDa) corresponding to the ER forms of GFP-GP4 (Bernat-Silvestre et al, 2020) and V-FLA11 (Supplemental Figure 8), respectively. In the *pgap1-3* mutant there was a strong decrease in the smear form (plasma membrane) of both proteins with a concomitant increase in their ER forms. **C-D.** Left panels (GFP-AGP4). Membrane fractions were obtained from PNSs of wild-type (**C**) or *pgap1-3* mutant (**D**) seedlings expressing GFP-AGP4 and incubated in the absence or presence of PI-PLC. Then, membranes were pelleted by centrifugation and pellets (P) and supernatants (S) were analyzed by SDS-PAGE and immunoblotting with antibodies against GFP to detect GFP-AGP4. Right panels (V-FLA11). PI-PLC treatment was performed directly in the PNSs, membranes were also pelleted by centrifugation and pellets and supernatants analyzed as before. In C, the upper part of the immunoblotting shows the smear form of GFP-AGP4 (left) or V-FLA11 (right) while the lower part highlights the ER bands of GFP-AGP4 (left) or V-FLA11 (right). Notice the decrease of both the smear forms and the lower ER bands from the pellet fraction and their partial appearance in the supernatant in wild-type seedlings. The presence of both forms in the supernatant seems to be only partial, probably due to degradation upon release from the membranes. In contrast, both forms are PI-PLC resistant in the *pgap1-3* mutant.

To test if the involvement of *PGAP1* in trafficking of GPI-APs was specific for this *PGAP*, we also tested the localization of GFP-AGP4, V-FLA11 and GFP-GPI in T-DNA insertion mutants of the PGAP1-like genes At2g44970, At3g52570 and At4g34310 (Supplemental Table 1), encoding putative GPI inositol-deacylase proteins but not expected to localize to compartments of the secretory pathway (Supplemental Figure 5). As shown in Supplemental Figure 6, these GPI-APs localized to the plasma membrane in these mutants, as it was the case of the transmembrane plasma membrane protein PIP2A-RFP. This suggests that these PGAP1-like genes encode proteins that are not required for transport from the ER to the plasma membrane of GPI-anchored proteins.

We next analyzed the localization of GFP-AGP4 and GFP-GPI by an alternative transient expression system, *Arabidopsis* protoplasts. In protoplasts from wild-type *Arabidopsis* plants, GFP-AGP4 and GFP-GPI mostly localized to the plasma membrane, as we have shown previously (Bernat-Silvestre et al., 2020). In contrast, GFP-AGP4 and GFP-GPI showed a predominant ER localization pattern in protoplasts from the *pgap1-1* mutant (Figure 7). The ER localization of GFP-AGP4 and GFP-GPI in *pgap1* mutant protoplasts was confirmed by colocalization experiments. As shown in Figure 7, both GFP-AGP4 and GFP-GPI strongly colocalized with two different ER marker proteins, RFP-calnexin and RFP-p24δ5. We could also detect the presence of both GFP-AGP4 and GFP-GPI at the plasma membrane, as shown by colocalization with Fei Mao styryl dye FM4-64, a lipid probe routinely used to label the plasma membrane (Figure 7). This suggests that a fraction of these GPI-APs can reach the plasma membrane in *pgap1* mutants, as observed also biochemically (see below, Figure 8). To test if the lack of PGAP1 enzymes affects the localization of other plasma membrane proteins different from GPI-APs, we used different membrane-anchoring types of minimal constructs, including a myristoylated and palmitoylated GFP (MAP-GFP) and a prenylated GFP (GFP-PAP) (Martinière et al., 2012). We also used a transmembrane protein, a GFP fusion with the plasma membrane ATPase (GFP-PMA). As shown in Supplemental Figure 7, these 3 proteins mainly localized to the plasma membrane of *pgap1-1* protoplasts, as in protoplasts from wild-type *Arabidopsis* plants. Therefore, PGAP1 function seems to be specifically required for ER export and transport to the plasma membrane of GPI-APs.

### 3. GPI anchor remodeling in *pgap1* mutants

We next used biochemical approaches to investigate the putative function of *Arabidopsis* PGAP1 in GPI anchor remodeling (in particular inositol diacylation) of GPI-APs. We have recently shown that GFP-AGP4 is a GPI-anchored protein (Bernat-Silvestre et al., 2020). Here, we have now biochemically characterized V-FLA11. To this end, a post-nuclear supernatant (PNS) from *N. benthamiana* leaves expressing V-FLA11 was analyzed by SDS-PAGE and immunoblotting with antibodies against GFP, to detect the Venus tag of V-FLA11 (Supplemental Figure 8). Similar to GFP-AGP4, two forms of V-FLA11 were detected, a predominant higher molecular weight smearing form (≈ 75 kDa), that may correspond to the mature form of the moderately glycosylated V-FLA11, and a much less abundant smaller molecular weight form (≈ 65 kDa) that may correspond to the ER form of V-FLA11 (Supplemental Figure 8A). To confirm this, leaves expressing V-FLA11 were treated with brefeldin A (BFA), to accumulate newly synthesized proteins at the ER. In the absence of BFA, V-FLA11 mainly localized to the plasma membrane, as expected (Supplemental Figure 8C). However, in the presence of BFA, V-FLA11 accumulated at the ER, where it showed a high degree of colocalization with the ER membrane protein RFP-p24δ5 (Supplemental Figure 8C). As shown in Supplemental Figure 8A, BFA treatment produced a drastic reduction of the 75 kDa smearing form and a concomitant increase in the 65 kDa band. This strongly suggests that 65-kD band corresponds to the ER form of V-FLA11, whereas the 75 kD smear should be the glycosylated form present at the Golgi/plasma membrane. In the case of GFP-AGP4 we also detected a 115 kDa smear form, which corresponds to the plasma membrane form, and a 70 kDa band which was shown to correspond to the ER form of GFP-AGP4 (Bernat-Silvestre et al., 2020).

To demonstrate that V-FLA11 is a GPI-anchored protein, we used Triton X-114 extraction and phosphatidylinositol-specific phospholipase C (PI-PLC) treatment. PNS from *N. benthamiana* leaves expressing V-FLA11 was extracted with Triton X-114 (TX114) and the TX114 detergent phase (containing V-FLA11) was treated or not with PI-PLC, as described in “Materials and Methods”. PI-PLC hydrolyzes the phosphodiester bond of the phosphatidylinositol, thereby releasing the protein from the membrane (Low, 1989). As shown in Supplemental Figure 8B, both the plasma membrane smear and the ER form of V-FLA11 were sensitive to PI-PLC, and moved from the detergent to the aqueous phase, thus confirming the GPI anchoring of V-FLA11.

To characterize GFP-AGP4 and V-FLA11 in wild-type and *pgap1 Arabidopsis* mutants, both proteins were transiently expressed in seedlings and PNSs were analyzed by SDS-PAGE and immunoblotting with GFP antibodies, as before. As shown in Figure 8, when both proteins were expressed in wild-type seedlings, we also detected a higher molecular weight smear form, which represents the plasma membrane form of both proteins, and a lower molecular weight band which should correspond to their ER forms. In the case of V-FLA11, the smear form had a higher molecular weight than that in *N. benthamiana* leaves. Remarkably, when both proteins were expressed in the *pgap1-3* mutant we found a drastic reduction of the higher molecular weight smear forms and a concomitant increase in the lower molecular weight bands (Figure 8). This is consistent with the predominant ER localization of both GFP-AGP4 and V-FLA11 in *pgap1* mutant shown before (Figures 5 and 6).

Mature GPI-APs are usually sensitive to PI-PLC because they have the 2-position of inositol free, leading to the release of the protein portions. In contrast, precursors of the GPI-anchor that contain an acyl chain linked to the 2-position of inositol are resistant to PI-PLC (Kinoshita and Fujita, 2016). Therefore, PI-PLC can be used as a tool to determine whether GPI-APs in the *pgap1* mutants contained or not the acyl chain in the inositol ring. Thus, GFP-AGP4 and V-FLA11 were transiently expressed in wild-type and *pgap1* mutant seedlings and post-nuclear supernatants were obtained. Membranes were pelleted by centrifugation, resuspended in buffer and treated or not with PI-PLC. After the treatment, membranes were pelleted again by centrifugation and pellets (membranes) and supernatants (containing released proteins) were analyzed by SDS-PAGE and immunoblotting with GFP antibodies. As shown in Figure 8C, in wild-type seedlings both the plasma membrane smear and the ER band of GFP-AGP4 and V-FLA11 were sensitive to PI-PLC, as shown before for GFP-AGP4 (Bernat-Silvestre et al., 2020) and for V-FLA11 in *N. benthamiana* (Supplemental Figure 8). In the *pgap1* mutant, the predominant form of both GFP-AGP4 and V-FLA11 is the lower molecular weight ER band. As shown in Figure 8D, neither GFP-AGP4 nor V-FLA11 were released from the membranes upon PI-PLC treatment. Resistance to PI-PLC treatment applied to both the main lower molecular weight ER band, but also to the small amount of the smear form, which corresponds to the fraction of these proteins reaching the plasma membrane. These results suggest that both GFP-AGP4 and V-FLA11 remain membrane attached in the *pgap1* mutant upon PI-PLC treatment, probably due to a defect in inositol deacylation.

Interestingly, after plasmolysis, most of the cell surface GFP-AGP4 was detected in the apoplast in wild-type seedlings, in contrast to the transmembrane plasma membrane protein SCAMP1-YFP, which mostly appears at the plasma membrane and in Hetchian strands (attachment sites between the plasma membrane and cell wall) upon mannitol treatment (Supplemental Figure 9). This is consistent with the fact that the highly glycosylated plasma membrane form of GFP-AGP4 (the 115-kD smear) had been shown to have a high tendency to appear in the aqueous phase after TX114 extraction (Bernat-Silvestre et al., 2020) (Supplemental Figure 9G). This localization is in agreement with the proposal that specific plasma membrane phospholipases may allow the release of some AGPs into the cell wall (Schultz et al., 1998; Ellis et al., 2010; Showalter and Basu, 2016; Ma et al., 2018). In *pgap1* mutants, SCAMP1-YFP was still found at the plasma membrane and Hetchian strands. Interestingly, the small fraction of GFP-AGP4 reaching the cell surface in *pgap1* mutants was found at the plasma membrane and Hetchian strands and was not released to the apoplast (Supplemental Figure 9B,C). This was confirmed biochemically by Triton X-114 extraction. Apart from the fact that the predominant form of GFP-AGP4 in *pgap1* mutants was the lower molecular weight ER band, which partitioned in the detergent phase upon Triton X-114 extraction, the plasma membrane smear of GFP-AGP4 also partitioned in the detergent phase, as the mitochondrial marker VDAC (Supplemental Figure 9H). This suggests that GFP-AGP4 remains membrane attached in *pgap1* mutants, probably due to the presence of the acyl group in the inositol moiety of the GPI anchor due to reduced PGAP1 activity.

## DISCUSSION

Most of the genes involved in GPI anchor assembly and their remodeling have orthologs in *Arabidopsis* (Luschnig and Seifert, 2011). However, it has to be established if all these orthologs are functional and if their function is conserved in plants. Five *Arabidopsis* orthologs of enzymes involved in the biosynthesis and attachment of the GPI anchor have been previously studied: SETH1, SETH2, PEANUT1 (PNT1), APTG1 and AtGPI8 (Lalanne et al., 2004; Gillmor et al., 2005; Dai et al., 2014; Bundy et al., 2016; Desnoyer et al., 2020). *Arabidopsis* null mutants of these enzymes are either gametophytic or embryogenic lethal mutants. This indicates that GPI-anchored proteins are essential for plant growth and development. However, no previous characterization of *Arabidopsis* GPI anchor lipid remodeling enzymes had been reported. In this study, we have undertaken the characterization of mutants of an *Arabidopsis* ortholog of the enzyme involved in the first step of lipid remodeling of the GPI anchor, yeast Bst1p/mammalian PGAP1.

Arabidopsis *PGAP1* (At3g27325) encodes a putative ortholog of yeast Bst1p/mammalian PGAP1, that are involved in inositol deacylation (Tanaka et al., 2004; Fujita et al., 2006b); (Luschnig and Seifert, 2011). We have found that Arabidopsis PGAP1 localizes in the ER and not at the Golgi apparatus or ER export sites. To investigate PGAP1 function, we have characterized loss of function mutants of *PGAP1*. In humans, null mutations of *PGAP1* are viable but result in defects in neuronal cell function (Ueda et al., 2007; Murakami et al., 2014; Williams et al., 2015). *ScBst1* is non-essential in yeast (Komath et al., 2018) and *ScBst1* mutants grew as well as wild-type at all temperatures (Elrod-Erickson and Kaiser, 1996; Fujita et al., 2006b). In *Candida Albicans*, deletion of *Bst1* impaired host infection and caused altered cell wall polysaccharides (Liu et al., 2016). Here, we show that knock-down *Arabidopsis pgap1* mutants displayed only mild phenotypes under standard growth conditions, but also showed an alteration in cell wall composition.

Interestingly, the trafficking of GPI-APs was altered in the *pgap1* mutants and they showed mainly ER localization in transient expression experiments both in protoplasts and seedlings. This agrees with previous results in yeast and mammals. Deletion of *ScBst1* caused delay in transport of GPI-APs from the ER to the Golgi and null mutations of PGAP1 in cultured cells caused accumulation of GPI-APs in the ER due to inefficient exit from the ER (Tanaka et al., 2004; Fujita et al., 2006b). However, steady state levels of cell surface GPI-APs were only mildly affected and GPI-APs were expressed at the cell surface, although with unusual GPI structures (three-“footed” GPI). We have also observed that a proportion GPI-APs could still reach the cell surface in *Arabidopsis pgap1* mutants, as in mammals and yeast. Over 40 % of the GPI-APs predicted by bioinformatics studies contain proteins with putative arabinogalactan (AG) or extensin-like glycosylation (Borner et al., 2003). Disrupted trafficking and/or glycosylation of GPI-APs could explain the altered composition of cell walls in *pgap1* mutants. Changes in Type II AG levels could result from reduced/altered AGP glycosylation and this potentially impacts cell wall assembly and architecture. AG glycans are proposed to cross-link to cell wall components, as is the case for ARABINOXYLAN PECTIN ARABINOGALACTAN PROTEIN1 (APAP1), shown to covalently cross-link to both pectins and arabinoxylans (Tan et al., 2013). Other GPI-APs can be involved in modulating both synthesis and remodelling of cell wall polymers, such as xyloglucan and cellulose. Members of the COBRA-like family of GPI-APs are known to regulate the deposition of cellulose into the wall (Li et al., 2013; Ben-Tov et al., 2015). A COBRA-like protein, BC1, has been shown to directly bind cellulose to affect MF crystallinity (Liu et al., 2013). Two putative GPI-anchored aspartic proteases, A36 and A39, were shown to co-localize with COBRA-LIKE 10 (Gao et al., 2017a; Gao et al., 2017b). In the apical cell walls of pollen tubes in double *a36a39* mutants, increased levels of highly methyl esterified homogalacturonan pectins and xyloglucans were detected. Changes in cell walls that compromise integrity, either during growth or as a result of damage from abiotic or biotic stresses, are detected by cell wall integrity sensors. GPI-APs are proposed to form part of the cell wall sensing complexes and compromised function could also contribute to changes in cell wall composition.

In the case of GFP-AGP4, we have found that in wild-type plants most of this arabinogalactan protein was found in the apoplast (and not at the plasma membrane), supporting the idea that AGP4 may be secreted into the extracellular matrix perhaps upon the action of plasma membrane phospholipases, as proposed for some of these proteins (Schultz et al., 1998; Ellis et al., 2010; Ma et al., 2018; Yeats et al., 2018). A similar behavior has been recently described for the arabinogalactan AGP21, which is also found at the plasma membrane and the apoplast (Borassi et al., 2020). In contrast, other AGPs remain attached to the plasma membrane, such as LeAGP1 (Sun et al., 2004), AGP17 (Sun et al., 2005; Zhang et al., 2011) and AGP18 (Yang and Showalter, 2007; Zhang et al., 2011). In animals, substrate-specific mechanisms have been proposed to release GPI-APs from the cell surface, including the phosphatidylinositol-specific phospholipases C and D (PI-PLCs and PI-PLDs) (Yeats et al., 2018). In contrast, no GPI-specific phospholipases have been yet characterized in plants. The identification of both specific and general GPI-cleavage mechanisms in plants may be an interesting area of future studies to understand the function of specific AGPs. The fact that GFP-AGP4 remained attached to the plasma membrane in *pgap1* mutants suggests a possible role of PGAP1 in secretion of certain GPI-APs, which could be important for signaling or cell wall localization of GPI-APs.

Finally, the ER localization of PGAP1-RFP and the PI-PLC resistance of GPI-APs in *pgap1* mutants suggest that PGAP1 indeed functions in *Arabidopsis* as the GPI inositol-deacylase which cleaves the acyl chain from the inositol ring of the GPI. Therefore, the results obtained indicate that PGAP1 is involved in GPI inositol deacylation and that inositol deacylation is important for efficient ER-to-Golgi transport of GPI-APs. To our knowledge, this is the first functional characterization of an enzyme involved in remodeling of the GPI anchor in plants. The characterization of other enzymes of the remodeling pathway should provide clues on the GPI anchor remodeling pathway in plants, to elucidate whether the pathway is completed at the ER, as in yeast, or if GPI anchor remodeling is completed in the Golgi apparatus (as in mammals). Future studies should also aim to elucidate more plant GPI anchor structures and the role of GPI anchor remodeling in sorting of GPI-APs into specific ER export sites and COPII vesicles or in their fate at the cell surface (plasma membrane versus apoplast or cell wall localization).

## METHODS

### Plant material

*Arabidopsis thaliana* (ecotype Col-0) was used as wild type. T-DNA insertion mutants used in this study were obtained from the Nottingham Arabidopsis Stock Centre. *A. thaliana* plants were grown in growth chambers as previously described (Ortiz-Masia et al., 2007). The T-DNA insertion mutants were characterized by PCR (Supplemental Table 3). Wild-type *Nicotiana benthamiana* plants were grown from surface-sterilized seeds on soil in the greenhouse at 24°C with 16 h daylength.

### RT-PCR

Total RNA was extracted from seedlings by using a Qiagen RNeasy plant mini kit, and 3 μg of the RNA solution was reverse-transcribed using the maxima first-strand cDNA synthesis kit for quantitative RT-PCR (Fermentas) according to the manufacturer’s instructions. Semi-quantitative PCRs (sqPCRs) were performed on 3 μl of cDNA template using Emerald Amp Max PCR Master Mix (Takara). The sequences of the primers used for PCR amplifications are included in Supplemental Table 4.

### Constructs and antibodies

The coding sequence of PGAP1-RFP, GFP-CESA3 and GFP-SEC24C were commercially synthesized de novo (Geneart AG) based on the sequence of the PGAP1 (At3g27325) and RFP, CESA3 (At5g05170) and GFP and SEC24C (At4g32640) and GFP, respectively and cloned into pCHF3 (Ortiz-Masia et al., 2007). A pGreenII0179 vector backbone (Hellens et al., 2000) was used for constructing V-FLA11 driven by pro35S: pGreen0179-35S-spFLA11-His-YFP-FLA11. The FLA11 (AT5G03170) coding sequence was amplified from *Arabidopsis* cDNA and assembled into a pGreen0179-35S-spFLA11-His-YFP vector using NEBuilder HiFi DNA Assembly kit (NEW ENGLAND Biolabs) according to the manufacturer’s instructions. Primers for amplification of FLA11 without a signal peptide are as follows: YMV101-FLA11-F, CAGGCGGAGGTGGGTCAcctaggCAGGCTCCAGCTCCAGGC; YMV101-FLA11-R, CATTAAAGCAGGACTCTAGATTATAT-CCACAGAGAAGAAGAAGCAG. The constructs used for transient expression experiments were: GFP-AGP4, GFP-GPI, MAP-GFP and GFP-PAP (Martinière et al., 2012; Bernat-Silvestre et al., 2020), GFP-PMA (Kim et al., 2001), PIP2A-RFP (Nelson et al., 2007) and RFP-calnexin (Künzl et al., 2016). Other constructs have been described previously: RFP-p24δ5 (Langhans et al., 2008; Montesinos et al., 2012); YFP-ManI (Nebenführ et al., 1999); GFP-HDEL (Pain et al., 2019); mCherry-HDEL (Nelson et al., 2007); OsSCAMP1-YFP (Lam et al., 2007), GFP-EMP12 (Gao et al., 2012), TIP1.1-GFP (Gattolin et al., 2011), SPΔCt-mCherry (Pereira et al., 2013). Antibodies against RFP and GFP were obtained from Clontech and Life Technologies, respectively.

### Transient gene expression in Arabidopsis protoplasts, Arabidopsis seedlings and *Nicotiana benthamiana* leaves

To obtain mesophyll protoplasts from Arabidopsis plants, the Tape-Arabidopsis Sandwich method was used, as described (Wu et al., 2009). Protoplasts were isolated from 4-week old rosette leaves. For transient expression, we used the PEG transformation method (Yoo et al., 2007). Transient expression of Arabidopsis seedlings by vacuum infiltration (Bernat-Silvestre et al., 2021) and *Nicotiana benthamiana* leaves mediated by *Agrobacterium tumefaciens* (Lerich et al., 2011) were performed as described previously.

### Preparation of protein extracts, PI-PLC treatment, SDS-PAGE and immunoblotting

*Nicotiana benthamiana* leaves or cotyledons from Arabidopsis seedlings expressing XFP-Proteins were frozen in liquid N2 and then grinded in homogenization buffer (HB, 0.3 M sucrose; 1 mM EDTA; 20 mM KCl; 20 mM HEPES pH 7.5), supplemented with 1mM DTT and a Protease Inhibitor Cocktail (Sigma), using a mortar and a pestle. The homogenate was centrifuged for 10 minutes at 1,200 xg and 4°C, and the post nuclear supernatant (PNS) was collected.

For treatment with Phosphatidylinositol-specific phospholipase C (PI-PLC) following transient expression of V-FLA11 in *Nicotiana benthamiana* leaves, PNS was incubated in the presence of 2 % TX-114 for 30 min at 4°C and then centrifuged 5 min at 16,000 g to pellet insoluble material. The supernatant was collected and incubated for 10 min at 37°C to achieve phase partitioning. The mixture was centrifuged 10 min at 20,000 g and 25 °C and the upper aqueous phase (A) and lower detergent phases (D) were collected. The detergent phase was diluted with TBS and incubated in the absence or presence of 2 U PI-PLC (from *Bacillus cereus*, 100 U/mL, Invitrogen^®^) for 1 h at 37°C. After this, samples were centrifuged again 10 min at 20,000 g and 25 °C to separate aqueous and detergent phases. Aqueous and detergent fractions were analyzed by SDS-PAGE and immunoblotting with GFP antibodies (to detect GFP-AGP4 and V-FLA11). For treatment with Phosphatidylinositol-specific phospholipase C (PI-PLC) following transient expression of GFP-AGP4 or V-FLA11 in Arabidopsis seedlings, a PNS was obtained. In the case of GFP-AGP4, membranes were pelleted by centrifugation of the PNS for 1 h at 150,000 xg. PNS (V-FLA11) or membrane fractions (GFP-AGP4) were then incubated in the absence or presence of PI-PLC, as before, and centrifuged for 1 h at 150,000 xg and 4°C, in order to separate membranes (pellet) from proteins released from the membranes upon PI-PLC treatment (supernatant). Membrane fractions were extracted with a lysis buffer containing 150 mM NaCl, 0.5 mM DTT, 0.5 % Triton X-100 and 50 mM Tris-HCl pH 7.5. After a 5 min centrifugation at 16,000 xg to remove detergent-insoluble material, membrane extracts were analyzed by SDS-PAGE and immunoblotting. Immunoblots were developed using the SuperSignal West Pico chemiluminescent substrate (Pierce,ThermoScientific) and analyzed using the ChemiDoc XRS + imagingsystem (Bio-Rad, http://www.bio-rad.com/). Immunoblots in the linear range of detection were quantified using Quantity One software (Bio-Rad Laboratories).

### Confocal Microscopy

Confocal fluorescent images were collected using an Olympus FV1000 confocal microscope with 603 water lens. Fluorescence signals for GFP (488 nm/496–518 nm), and RFP (543 nm/593–636 nm) were detected. Sequential scanning was used to avoid any interference between fluorescence channels. Post-acquisition image processing was performed using the FV10-ASW 4.2 Viewer and ImageJ (v.1.45).

### Cell wall linkage analysis

De-starched AIR samples were prepared from 7 day old seedlings grown on MS media with 1% sucrose. AIR was carboxyl reduced and methylated for linkage analysis according to the method outlined in Pettolino et al., (2012). The resulting permethylated alditol acetates were separated and quantified by GC-MS as described in Pettolino et al., (2012). Polysaccharide composition was deduced from the linkage analyses. Two biological replicates with two technical replicates each were measured. Data shown as average.

## ACKNOWLEDGEMENTS

F.A. was supported by the Ministerio de Economía y Competitividad (grant no BFU2016-76607-P) and Generalitat Valenciana (ACOMP/2014/202 and AICO/2020/187 to F.A. and M.J.M.). C.B.S. and J.S.S. were recipient of a fellowship from Ministerio de Ciencia, Innovación y Universidades (FPU program). C.B.S. was recipient of an EMBO short-term fellowship and a short-term fellowship from Ministerio de Ciencia, Innovación y Universidades. We thank John Runions for the GFP-AGP4, GFP-GPI, MAP-GFP and GFP-PAP constructs, Inhwan Hwang for the GFP-PMA construct, Liwen Jiang for the GFP-EMP12 and OsSCAMP1-YFP constructs, Lorenzo Frigerio for the TIP1.1-GFP construct and Claudia Pereira for the SPΔCt-mCherry construct. We thank the microscopy section and the greenhouse of SCSIE (University of Valencia).

## REFERENCES

Ben-Tov D, Abraham Y, Stav S, Thompson K, Loraine A, Elbaum R, de Souza A, Pauly M, Kieber JJ, Harpaz-Saad S (2015) COBRA-LIKE2, a Member of the Glycosylphosphatidylinositol-Anchored COBRA-LIKE Family, Plays a Role in Cellulose Deposition in Arabidopsis Seed Coat Mucilage Secretory Cells. Plant Physiol 167: 711–724

Bernat-Silvestre C, De Sousa Vieira V, Sánchez-Simarro J, Aniento F, Marcote MJ (2021) Transient Transformation of A. thaliana Seedlings by Vacuum Infiltration. Methods Mol. Biol. Humana Press Inc., pp 147–155

Bernat-Silvestre C, Vieira VDS, Sanchez-Simarro J, Pastor-Cantizano N, Hawes C, Marcote MJ, Aniento F (2020) P24 Family proteins are involved in transport to the plasma membrane of GPI-anchored proteins in plants. Plant Physiol 184: 1333–1347

Borassi C, Gloazzo Dorosz J, Ricardi MM, Carignani Sardoy M, Pol Fachin L, Marzol E, Mangano S, Rodríguez Garcia DR, Martínez Pacheco J, Rondón Guerrero Y del C, et al (2020) A cell surface arabinogalactan-peptide influences root hair cell fate. New Phytol 227: 732–743

Borner GHH, Lilley KS, Stevens TJ, Dupree P (2003) Identification of glycosylphosphatidylinositol-anchored proteins in Arabidopsis. A proteomic and genomic analysis. Plant Physiol 132: 568–577

Bosson R, Jaquenoud M, Conzelmann A (2006) GUP1 of Saccharomyces cerevisiae encodes an O-acyltransferase involved in remodeling of the GPI anchor. Mol Biol Cell 17: 2636–45

Bundy MGR, Kosentka PZ, Willet AH, Zhang L, Miller E, Shpak ED (2016) A mutation in the catalytic subunit of the glycosylphosphatidylinositol transamidase disrupts growth, fertility, and stomata formation. Plant Physiol 171: 974–985

Castillon GA, Aguilera-Romero A, Manzano-Lopez J, Epstein S, Kajiwara K, Funato K, Watanabe R, Riezman H, Muñiz M (2011) The yeast p24 complex regulates GPI-anchored protein transport and quality control by monitoring anchor remodeling. Mol Biol Cell 22: 2924–2936

Castillon GA, Watanabe R, Taylor M, Schwabe TME, Riezman H (2009) Concentration of GPI-anchored proteins upon ER exit in yeast. Traffic 10: 186–200

Cheung AY, Li C, Zou YJ, Wu HM (2014) Glycosylphosphatidylinositol anchoring: Control through modification. Plant Physiol 166: 748–750

Dai XR, Gao X-Q, Chen GH, Tang LL, Wang H, Zhang XS (2014) ABNORMAL POLLEN TUBE GUIDANCE1, an Endoplasmic Reticulum-Localized Mannosyltransferase Homolog of GLYCOSYLPHOSPHATIDYLINOSITOL10 in Yeast and PHOSPHATIDYLINOSITOL GLYCAN ANCHOR BIOSYNTHESIS B in Human, Is Required for Arabidopsis Pollen Tube Micropylar Gu. Plant Physiol 165: 1544–1556

Desnoyer N, Howard G, Jong E, Palanivelu R (2020) AtPIG-S, a predicted Glycosylphosphatidylinositol Transamidase subunit, is critical for pollen tube growth in Arabidopsis. BMC Plant Biol. doi: 10.1186/s12870-020-02587-x

Ellis M, Egelund J, Schultz CJ, Bacic A (2010) Arabinogalactan-proteins: Key regulators at the cell surface? Plant Physiol 153: 403–419

Elrod-Erickson MJ, Kaiser CA (1996) Genes that control the fidelity of endoplasmic reticulum to Golgi transport identified as suppressors of vesicle budding mutations. Mol Biol Cell 7: 1043–1058

Finn RD, Mistry J, Tate J, Coggill P, Heger A, Pollington JE, Gavin OL, Gunasekaran P, Ceric G, Forslund K, et al (2010) The Pfam protein families database. Nucleic Acids Res 38: D211–22

Fujita M, Umemura M, Yoko-o T, Jigami Y (2006a) PER1 is required for GPI-phospholipase A2 activity and involved in lipid remodeling of GPI-anchored proteins. Mol Biol Cell 17: 5253–64

Fujita M, Yoko-O T, Jigami Y (2006b) Inositol deacylation by Bst1p is required for the quality control of glycosylphosphatidylinositol-anchored proteins. Mol Biol Cell 17: 834–850

Gao C, Yu CKY, Qu S, San MWY, Li KY, Lo SW, Jiang L (2012) The Golgi-Localized Arabidopsis Endomembrane Protein12 Contains Both Endoplasmic Reticulum Export and Golgi Retention Signals at Its C Terminus. Plant Cell 24: 2086–2104

Gao H, Li R, Guo Y (2017a) Arabidopsis aspartic proteases A36 and A39 play roles in plant reproduction. Plant Signal Behav 12: 219–239

Gao H, Zhang Y, Wang W, Zhao K, Liu C, Bai L, Li R, Guo Y (2017b) Two Membrane-Anchored Aspartic Proteases Contribute to Pollen and Ovule Development. Plant Physiol 173: 219–239

Gattolin S, Sorieul M, Frigerio L (2011) Mapping of tonoplast intrinsic proteins in maturing and germinating Arabidopsis seeds reveals dual localization of embryonic TIPs to the tonoplast and plasma membrane. Mol Plant 4: 180–189

Ghugtyal V, Vionnet C, Roubaty C, Conzelmann A (2007) CWH43 is required for the introduction of ceramides into GPI anchors in Saccharomyces cerevisiae. Mol Microbiol 65: 1493–1502

Gillmor CS, Lukowitz W, Brininstool G, Sedbrook JC, Hamann T, Poindexter P, Somerville C (2005) Glycosylphosphatidylinositol-Anchored Proteins Are Required for Cell Wall Synthesis and Morphogenesis in Arabidopsis. Plant Cell 17:1128–1140

Hellens RP, Anne Edwards E, Leyland NR, Bean S, Mullineaux PM (2000) pGreen: A versatile and flexible binary Ti vector for Agrobacterium-mediated plant transformation. Plant Mol Biol 42: 819–832

Hemsley PA (2015) The importance of lipid modified proteins in plants. New Phytol 205: 476–489

Hofmann K, Stoffel W (1993) TMbase - A database of membrane spanning proteins segments.

Hruz T, Laule O, Szabo G, Wessendorp F, Bleuler S, Oertle L, Widmayer P, Gruissem W, Zimmermann P (2008) Genevestigator V3: A Reference Expression Database for the Meta-Analysis of Transcriptomes. Adv Bioinformatics 2008: 1–5

Hunter S, Apweiler R, Attwood TK, Bairoch A, Bateman A, Binns D, Bork P, Das U, Daugherty L, Duquenne L, et al (2009) InterPro: The integrative protein signature database. Nucleic Acids Res. doi: 10.1093/nar/gkn785

Johnson KL (2003) The Fasciclin-Like Arabinogalactan Proteins of Arabidopsis. A Multigene Family of Putative Cell Adhesion Molecules. Plant Physiol 133: 1911–1925

Kim DH, Eu Y-J, Yoo CM, Kim Y-W, Pih KT, Jin JB, Kim SJ, Stenmark H, Hwang I (2001) Trafficking of phosphatidylinositol 3-phosphate from the trans - Golgi network to the lumen of the central vacuole in plant cells. Plant Cell 13: 287–301

Kinoshita T (2020) Biosynthesis and biology of mammalian GPI-anchored proteins. Open Biol. doi: 10.1098/rsob.190290

Kinoshita T, Fujita M (2016) Biosynthesis of GPI-anchored proteins: special emphasis on GPI lipid remodeling. J Lipid Res 57: 6–24

Komath SS, Singh SL, Pratyusha VA, Sah SK (2018) Generating anchors only to lose them: The unusual story of glycosylphosphatidylinositol anchor biosynthesis and remodeling in yeast and fungi. IUBMB Life 70: 355–383

Künzl F, Früholz S, Fäßler F, Li B, Pimpl P (2016) Receptor-mediated sorting of soluble vacuolar proteins ends at the trans-Golgi network/early endosome. Nat Plants 2: 16017

Lalanne E, Honys D, Johnson A, Borner GHH, Lilley KS, Dupree P, Grossniklaus U, Twell D (2004) SETH1 and SETH2, two components of the glycosylphosphatidylinositol anchor biosynthetic pathway, are required for pollen germination and tube growth in Arabidopsis. Plant Cell 16: 229–40

Lam SK, Siu CL, Hillmer S, Jang S, An G, Robinson DG, Jiang L (2007) Rice SCAMP1 Defines Clathrin-Coated, trans -Golgi–Located Tubular-Vesicular Structures as an Early Endosome in Tobacco BY-2 Cells. Plant Cell 19: 296–319

Langhans M, Marcote MJ, Pimpl P, Virgili-López G, Robinson DG, Aniento F (2008) In vivo Trafficking and Localization of p24 Proteins in Plant Cells. Traffic 9: 770–785

Lerich A, Langhans M, Sturm S, Robinson DG (2011) Is the 6 kDa tobacco etch viral protein a bona fide ERES marker? J Exp Bot 62: 5013–5023

Li S, Ge FR, Xu M, Zhao XY, Huang GQ, Zhou LZ, Wang JG, Kombrink A, McCormick S, Zhang XS, et al (2013) Arabidopsis COBRA-LIKE 10, a GPI-anchored protein, mediates directional growth of pollen tubes. Plant J 74: 486–497

Liu L, Shang-Guan K, Zhang B, Liu X, Yan M, Zhang L, Shi Y, Zhang M, Qian Q, Li J, et al (2013) Brittle Culm1, a COBRA-Like Protein, Functions in Cellulose Assembly through Binding Cellulose Microfibrils. PLoS Genet 9: e1003704

Liu W, Zou Z, Huang X, Shen H, He LJ, Chen SM, Li LP, Yan L, Zhang SQ, Zhang JD, et al (2016) Bst1 is required for Candida albicans infecting host via facilitating cell wall anchorage of Glycosylphosphatidyl inositol anchored proteins. Sci Rep. doi: 10.1038/srep34854

Low MG (1989) The glycosyl-phosphatidylinositol anchor of membrane proteins. BBA - Rev Biomembr 988: 427–454

Luschnig C, Seifert GJ (2011) Posttranslational modifications of plasma membrane proteins and their implications for plant growth and development. Plant Cell Monogr 19: 109–128

Ma Y, Zeng W, Bacic A, Johnson K (2018) AGPs Through Time and Space. Annu. Plant Rev. online. Wiley, pp 767–804

MacMillan CP, Mansfield SD, Stachurski ZH, Evans R, Southerton SG (2010) Fasciclin-like arabinogalactan proteins: specialization for stem biomechanics and cell wall architecture in Arabidopsis and Eucalyptus. Plant J 62: 689–703

Maeda Y, Tashima Y, Yoko-o T, Jigami Y, Fujita M, Kinoshita T, Houjou T, Taguchi R (2007) Fatty Acid Remodeling of GPI-anchored Proteins Is Required for Their Raft Association. Mol Biol Cell 18: 1497–1506

Manzano-Lopez J, Perez-Linero AM, Aguilera-Romero A, Martin ME, Okano T, Silva DV, Seeberger PH, Riezman H, Funato K, Goder V, et al (2015) COPII coat composition is actively regulated by luminal cargo maturation. Curr Biol 25: 152–162

Martinière A, Lavagi I, Nageswaran G, Rolfe DJ, Maneta-Peyret L, Luu D-T, Botchway SW, Webb SED, Mongrand S, Maurel C, et al (2012) Cell wall constrains lateral diffusion of plant plasma-membrane proteins. Proc Natl Acad Sci U S A 109:12805–10

Montesinos JC, Sturm S, Langhans M, Hillmer S, Marcote MJ, Robinson DG, Aniento F (2012) Coupled transport of Arabidopsis p24 proteins at the ER-Golgi interface. J Exp Bot 63: 4243–4261

Muñiz M, Riezman H (2016) Trafficking of glycosylphosphatidylinositol anchored proteins from the endoplasmic reticulum to the cell surface. J Lipid Res 57: 352–360

Muniz M, Zurzolo C (2014) Sorting of GPI-anchored proteins from yeast to mammals - common pathways at different sites? J Cell Sci 127: 2793–2801

Murakami Y, Tawamie H, Maeda Y, Büttner C, Buchert R, Radwan F, Schaffer S, Sticht H, Aigner M, Reis A, et al (2014) Null Mutation in PGAP1 Impairing Gpi-Anchor Maturation in Patients with Intellectual Disability and Encephalopathy. PLoS Genet 10: e1004320

Nebenführ A, Gallagher LA, Dunahay TG, Frohlick JA, Mazurkiewicz AM, Meehl JB, Staehelin LA (1999) Stop-and-Go Movements of Plant Golgi Stacks Are Mediated by the Acto-Myosin System. Plant Physiol 121: 1127–1141

Nelson BK, Cai X, Nebenführ A (2007) A multicolored set of in vivo organelle markers for co-localization studies in Arabidopsis and other plants. Plant J 51: 1126–1136

Ortiz-Masia D, Perez-Amador MA, Carbonell J, Marcote MJ (2007) Diverse stress signals activate the C1 subgroup MAP kinases of *Arabidopsis*. FEBS Lett 581: 1834–1840

Oxley D, Bacic A (1999) Structure of the glycosylphosphatidylinositol anchor of an arabinogalactan protein from Pyrus communis suspension-cultured cells. Proc Natl Acad Sci U S A 96: 14246–51

Pain C, Kriechbaumer V, Kittelmann M, Hawes C, Fricker M (2019) Quantitative analysis of plant ER architecture and dynamics. Nat Commun 10: 984

Pereira AM, Lopes AL, Coimbra S (2016) JAGGER, an AGP essential for persistent synergid degeneration and polytubey block in Arabidopsis. Plant Signal Behav 11: e1209616

Pereira C, Pereira S, Satiat-Jeunemaitre B, Pissarra J (2013) Cardosin A contains two vacuolar sorting signals using different vacuolar routes in tobacco epidermal cells. Plant J 76: 87–100

Pettolino FA, Walsh C, Fincher GB, Bacic A (2012) Determining the polysaccharide composition of plant cell walls. Nat Protoc 7: 1590–1607

Pittet M, Conzelmann A (2007) Biosynthesis and function of GPI proteins in the yeast Saccharomyces cerevisiae. Biochim Biophys Acta - Mol Cell Biol Lipids 1771: 405–420

Rodriguez-Gallardo S, Kurokawa K, Sabido-Bozo S, Cortes-Gomez A, Ikeda A, Zoni V, Aguilera-Romero A, Perez-Linero AM, Lopez S, Waga M, et al (2020) Ceramide chain length-dependent protein sorting into selective endoplasmic reticulum exit sites. Sci Adv. doi: 10.1126/sciadv.aba8237

Schultz C, Gilson P, Oxley D, Youl J, Bacic A (1998) GPI-anchors on arabinogalactan-proteins: Implications for signalling in plants. Trends Plant Sci 3: 426–431

Showalter AM, Basu D (2016) Glycosylation of arabinogalactan-proteins essential for development in Arabidopsis. Commun Integr Biol 9: e1177687

Silva L, De Almeida RFM, Fedorov A, Matos APA, Prieto M (2006) Ceramide-platform formation and -induced biophysical changes in a fluid phospholipid membrane. Mol Membr Biol 23: 137–148

Strasser R, Seifert G, Doblin MS, Johnson KL, Ruprecht C, Pfrengle F, Bacic A, Estevez JM (2021) Cracking the “Sugar Code”: A Snapshot of N- and O-Glycosylation Pathways and Functions in Plants Cells. Front Plant Sci 12: 157

Sun W, Xu J, Yang J, Kieliszewski MJ, Showalter AM (2005) The lysine-rich arabinogalactan-protein subfamily in Arabidopsis: Gene expression, glycoprotein purification and biochemical characterization. Plant Cell Physiol 46: 975–984

Sun W, Zhao ZD, Hare MC, Kieliszewski MJ, Showalter AM (2004) Tomato LeAGP-1 is a plasma membrane-bound, glycosylphosphatidylinositol-anchored arabinogalactan-protein. Physiol Plant 120: 319–327

Tan L, Eberhard S, Pattathil S, Warder C, Glushka J, Yuan C, Hao Z, Zhu X, Avci U, Miller JS, et al (2013) An Arabidopsis cell wall proteoglycan consists of pectin and arabinoxylan covalently linked to an arabinogalactan protein. Plant Cell 25: 270–87

Tanaka S, Maeda Y, Tashima Y, Kinoshita T (2004) Inositol Deacylation of Glycosylphosphatidylinositol-anchored Proteins Is Mediated by Mammalian PGAP1 and Yeast Bst1p. J Biol Chem 279: 14256–14263

Ueda Y, Yamaguchi R, Ikawa M, Okabe M, Morii E, Maeda Y, Kinoshita T (2007) PGAP1 knock-out mice show otocephaly and male infertility. J Biol Chem 282: 30373–30380

Umemura M, Fujita M, Yoko-O T, Fukamizu A, Jigami Y (2007) Saccharomyces cerevisiae CWH43 is involved in the remodeling of the lipid moiety of GPI anchors to ceramides. Mol Biol Cell 18: 4304–16

Williams C, Jiang YH, Shashi V, Crimian R, Schoch K, Harper A, McHale D, Goldstein D, Petrovski S (2015) Additional evidence that PGAP1 loss of function causes autosomal recessive global developmental delay and encephalopathy. Clin Genet 88: 597–599

Wu F-H, Shen S-C, Lee L-Y, Lee S-H, Chan M-T, Lin C-S (2009) Tape-Arabidopsis Sandwich - a simpler Arabidopsis protoplast isolation method. Plant Methods 5: 16

Yang J, Showalter AM (2007) Expression and localization of AtAGP18, a lysine-rich arabinogalactan-protein in Arabidopsis. Planta 226: 169–179

Yeats TH, Bacic A, Johnson KL (2018) Plant glycosylphosphatidylinositol anchored proteins at the plasma membrane-cell wall nexus. J Integr Plant Biol 60: 649–669

Yoko-o T, Umemura M, Komatsuzaki A, Ikeda K, Ichikawa D, Takase K, Kanzawa N, Saito K, Kinoshita T, Taguchi R, et al (2018) Lipid moiety of glycosylphosphatidylinositol-anchored proteins contributes to the determination of their final destination in yeast. Genes to Cells 23: 880–892

Yoo SD, Cho YH, Sheen J (2007) Arabidopsis mesophyll protoplasts: A versatile cell system for transient gene expression analysis. Nat Protoc 2: 1565–1572

Zhang Y, Yang J, Showalter AM (2011) AtAGP18 is localized at the plasma membrane and functions in plant growth and development. Planta 233: 675–683

Zimmermann P, Hirsch-Hoffmann M, Hennig L, Gruissem W (2004) GENEVESTIGATOR. Arabidopsis microarray database and analysis toolbox. Plant Physiol 136: 2621–2632

